# Mental individuation of imagined finger movements can be achieved using TMS-based neurofeedback

**DOI:** 10.1101/2021.02.23.432532

**Authors:** Ernest Mihelj, Marc Bächinger, Sanne Kikkert, Kathy Ruddy, Nicole Wenderoth

**Author notes:** **CORRESPONDING AUTHOR** Prof. Nicole Wenderoth, Neural Control of Movement Lab, Department of Health Sciences and Technology, ETH Zurich, Switzerland, Auguste-Piccard-Hof 1 Building HPT, Floor EETH Zurich, Switzerland. **HIGHLIGHTS** - TMS-based neurofeedback reinforces finger-selective modulation of corticomotor excitability - TMS-based neurofeedback causes strong beta desynchronization during motor imagery - EEG finger representation patterns became more separable following TMS-based neurofeedback.

## Abstract

Neurofeedback (NF) in combination with motor imagery (MI) can be used for training individuals to volitionally modulate sensorimotor activity without producing overt movements. However, until now, NF methods were of limited utility for mentally training specific hand and finger actions. Here we employed a novel transcranial magnetic stimulation (TMS) based protocol to probe and detect MI-induced motor activity patterns in the primary motor cortex (M1) with the aim to reinforce selective facilitation of single finger representations. We showed that TMS-NF training but not MI training with uninformative feedback enabled participants to selectively upregulate corticomotor excitability of one finger, while simultaneously downregulating excitability of other finger representations within the same hand. Successful finger individuation during MI was accompanied by strong desynchronisation of sensorimotor brain rhythms, particularly in the beta band, as measured by electroencephalography. Additionally, informative TMS-NF promoted more dissociable EEG activation patterns underlying single finger MI, when compared to MI of the control group where no such feedback was provided. Our findings suggest that selective TMS-NF is a new approach for acquiring the ability of finger individuation even if no overt movements are performed. This might offer new treatment modality for rehabilitation after stroke or spinal cord injury.

## INTRODUCTION

In recent years neurofeedback (NF) approaches have gained traction both in scientific and commercial domains. Due to its simplicity and high temporal resolution, electroencephalography (EEG) has emerged as the most popular neuroimaging modality used in non-invasive NF (Ramadan and Vasilakos, 2017). EEG based NF approaches generally extract intentions from brain states and translate these either into control commands that operate external devices, or into a form of neurofeedback that is fed back to the user (Birbaumer, 2006; Schwartz, 2004; Schwartz et al., 2006; Wolpaw et al., 2002). Interestingly, NF can be utilised to train participants to modulate their neural activity even in the absence of overt motor output (Enriquez-Geppert et al., 2017).

For the sensorimotor system, this is generally achieved via motor imagery (MI), i.e. participants mentally simulate movements. MI is most notably associated with the desynchronisation of sensorimotor rhythms (SMR), i.e. reduced oscillatory activity in the mu and beta frequency bands. MI induced changes in SMRs can be reliably measured using EEG in healthy participants but also in patients with brain injuries (e.g. stroke patients) or suffering from physical disability (e.g. paralyzed patients). Even though these patients struggle to generate overt movements, numerous studies have confirmed that they can modulate their SMRs by using MI and that MI-based training has beneficial effects on the restoration of impaired motor function (Barclay et al., 2020; Cramer et al., 2007; García Carrasco and Aboitiz Cantalapiedra, 2016; Guerra et al., 2017; Sun et al., 2013).

Research in healthy participants has shown that the effectiveness of MI training is enhanced when NF is provided to the user (Bai et al., 2014; Ono et al., 2014). However, when MI training involves more than one MI task, EEG based NF approaches are challenged because the temporal and spatial patterns of MI-induced SMR modulation are highly overlapping, making it difficult to distinguish between the different MI tasks. This challenge can be overcome by machine learning methods when the different MI tasks involve effectors with clearly distinct cortical representations, for example imagining left hand movements versus right hand movements which activate different hemispheres (Chu et al., 2018; Edelman et al., 2016; Tavakolan et al., 2017; Yong and Menon, 2015). By contrast, EEG-based NF is of limited utility when more fine-grained MI tasks need to be distinguished, like single finger movements (Hayashi et al., 2017), since the relatively low spatial resolution of EEG makes it difficult to reliably dissociate the generated brain activity patterns.

Here we investigate an alternative approach for supporting MI of single finger movements by NF and present a new protocol which uses transcranial magnetic stimulation (TMS) to detect MI-induced motor activity patterns in primary motor cortex (M1). The basic mechanism is that MI modulates the excitability of motor cortex when probed by TMS (Grosprêtre et al., 2016), a phenomenon which has previously been used to train participants to volitionally excite or inhibit corticomotor excitability (Kasai et al., 1997; Koganemaru et al., 2018; Majid et al., 2015; Ruddy et al., 2018). However, it is currently unclear whether participants can use TMS-NF to gain selective control of corticomotor excitability of single finger representations, which would be similar to finger individuation training as used in clinical settings (Xu et al., 2017). To answer this question, we developed a TMS-based NF (TMS-NF) approach where participants were trained to selectively facilitate corticomotor excitability of one finger while simultaneously inhibiting the other fingers. We recorded concurrent EEG in order to investigate how this type of TMS-NF training affects neural activity across the brain. It is well known that corticomotor excitability is closely related to rhythmic brain oscillatory activity (Jensen et al., 2011) such that higher corticomotor excitability is associated with stronger desynchronisation of SMRs (Zaehle et al., 2010). Accordingly, we hypothesized that TMS-NF will strongly affect mu and beta rhythms in electrodes overlying sensorimotor areas. We further hypothesize that repeated TMS-NF training would lead to increased separability of SMR related neural patterns underlying different types of motor imagery.

## MATERIALS AND METHODS

### Participants

Twenty volunteers participated in this experiment. Participants were randomly split into 2 groups: an experimental group (10 participants, age 26.2 ± 3.79, 3 females) and a control group (10 participants, age 25.4 ± 2.72, 4 females) which were highly similar in age and gender distribution. A power analysis was conducted a-priori to estimate the required sample size (see Supplementary figure 1 and Supplementary table 1). Based on the alpha value of 0.05, inclusion of two groups and an effect size of 1.27 (effect size reported in Ruddy et al., 2018) we estimated a sample size of n=10 per group. All participants were right-handed according to the Edinburgh Handedness Inventory (Oldfield, 1971) and had no TMS contraindications (Rossi et al., 2009; Wassermann, 1998). Upon study conclusion, participants were debriefed and financially compensated for their time and effort. None of the participants reported any major side effects resulting from the stimulation. Ethical approval was granted by the Kantonale Ethikkommission Zürich (KEK-ZH-Nr. 2018-01078) and written informed consent was obtained from all participants prior to study onset.

### Experimental protocol

The experiment consisted of five sessions of TMS-based neurofeedback training during which EEG was simultaneously recorded. Participants were instructed to selectively perform kinaesthetic imagery of the right thumb, index, or little finger (Figure 1).

**Figure 1.**
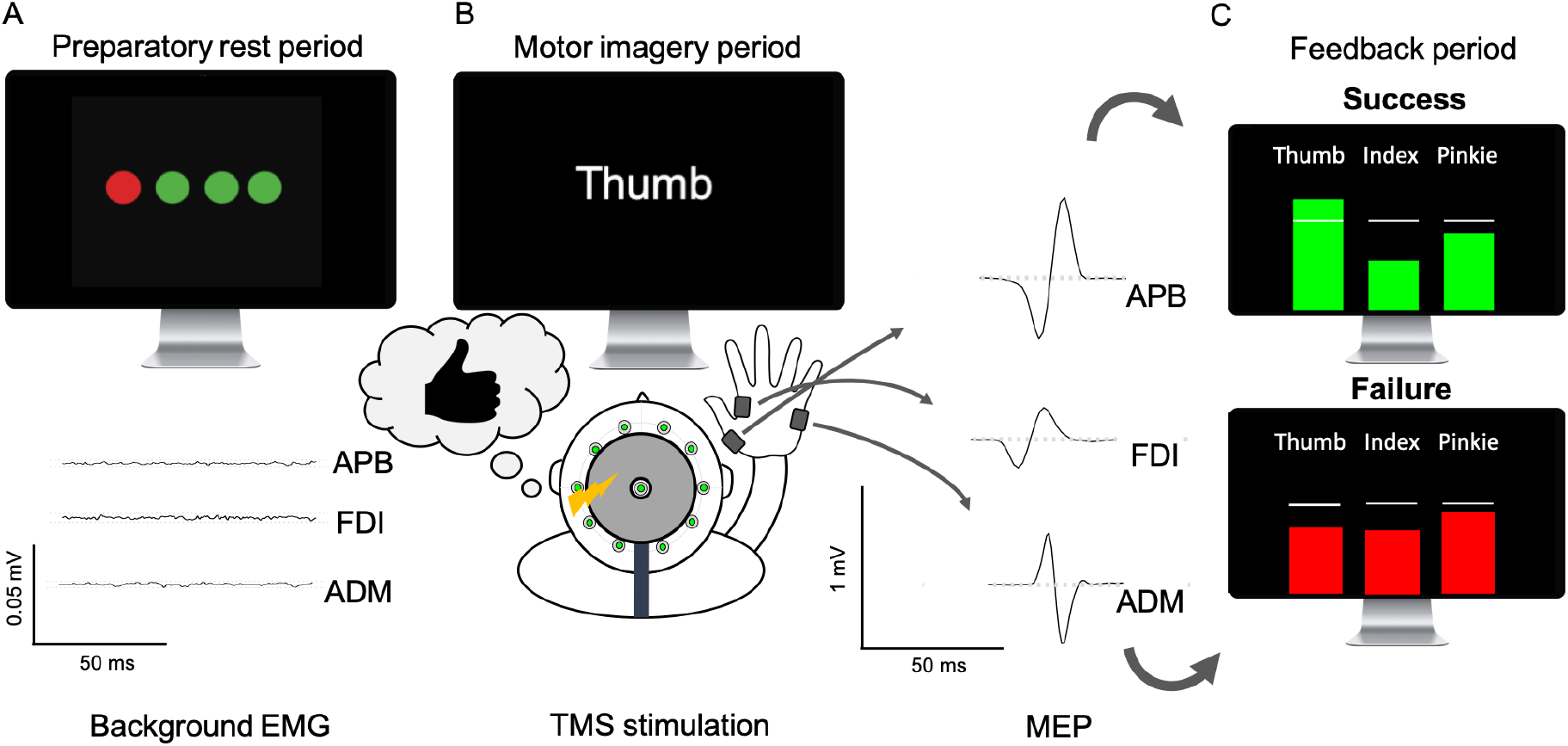
Experimental setup. A) Each trial started with a display of four circles representing the background EMG in the recorded hand muscles (circles from left to right = right APB, right FDI, right ADM, and left FDI). A circle turned red if the rest root mean squared (rms) EMG signal of that muscle was greater than seven microvolts. A trial would only proceed if all four circles were green for at least 500ms. B) When this requirement was fulfilled, a cue appeared indicating the start of the MI period that lasted for an interval jittered between 4-6 s. The participant was instructed to perform MI with the cued finger (thumb, index, or pinkie) until the TMS pulse was discharged. Also, during the MI period, it remained essential to keep the background EMG below 7 μV for a TMS pulse to be delivered. C) A TMS pulse was delivered 4-6 s after MI onset. The peak-to-peak amplitude of the motor evoked potentials (MEP) evoked by the TMS was calculated in real-time for the 3 muscles of the right hand. Each muscle’s MEP was divided by the average MEP value of the preceding 10 rest trials. The resulting normalised MEP values were presented as 3 bars (one for each muscle) to the participant in the feedback stage of the trial. The white horizontal lines indicate no change from the baseline for the respective muscle, i.e. normalized MEP value of 1. Subject were instructed that the bar of the cued target muscle should exceed the white line while the bars of the two other non-traget muscles should stay below. As long as neither of the normalised non-target muscles MEPs rose above 130 % of their respective baseline, and the target muscle MEP exceeded its baseline, a trial was deemed successful and the bars turned green. Otherwise they turned red indicated an unsuccessful trial.

Sessions were performed on five separate days. In sessions 1-3, MI trials were in distinct blocks for each finger (“blocked sessions”) while in sessions 4-5 MI trials of the different fingers were pseudo-randomly distributed (“interleaved sessions“). This training regime was chosen, because pilot work has shown that imagining the same finger movement several trials in a row is easier than switching to a new finger after every trial. Thus, over the 5 sessions, task difficulty increased for the last two sessions.

For the blocked sessions 1-3, participants underwent 42 MI trials and 20 rest trials for each finger which followed a fixed order (e.g. 10 rest – 21 MI thumb – 10 rest – 21 MI index −10 rest – 21 MI pinkie and so forth). After each block there was a short break. The order of fingers activated via MI was randomized. The interleaved sessions 4-5 followed the same principle structure but which finger movement to be imagined varied randomly within each set of 21 MI trails (10 rest – 21 MI random finger – 10 rest – 21 MI random finger).

Every MI trial consisted of a (i) preparatory rest period where subjects had to relax and were instructed which finger movement had to be imagined; (ii) MI period where participants imagined movements with the instructed finger; and (iii) feedback period where participants saw the modulation pattern for all three muscles and received some reinforcing feedback (see further details below). Every rest trial followed the same structure but participants were instructed to not engage in any MI.

Several possible MI strategies (Supplementary table 2) were provided to the participants prior to study onset, but they were instructed to explore their own optimal MI strategy with the aid of the NF.

### TMS procedures

Single-pulse monophasic TMS was delivered using a round coil with a 90 mm loop diameter which was connected to a Magstim 200 stimulator (Magstim, Whitland, UK). This coil generates a spherical magnetic field perpendicular to the coil which delivers less focal and more distributed stimulation, compared to the figure eight coil, making it suitable for protocols that target multiple finger representations. The coil was positioned over the vertex with the annulus under the coil approximately passing through the right and left primary motor hand area (M1; Groppa et al. 2012). The current direction was oriented to target left M1. The TMS coil was placed on a 3D-printed custom plastic coil spacer (Ruddy et al., 2018), which increases the distance between the coil and the head by 9mm. The coil spacer reduces artefacts relating to physical contact between the coil and the underlying EEG electrodes but it renders resting motor thresholds (RMTs) of all muscles slightly elevated (which was, however, not problematic for the current study).

Surface electromyography (EMG; Trigno Wireless; Delsys) was recorded from the left and right First Dorsal Interosseous (FDI), right Abductor Pollicis Brevis (APB) and right Abductor Digiti Minimi (ADM). The left FDI was included as a control muscle to detect whether participants might use a strategy whereby they activate left-hand muscles. EMG data were sampled at 2000 Hz (National Instruments, Austin, Texas), amplified and stored on a local PC for further analysis.

RMT was established as the lowest stimulation intensity that generated MEPs with a peak-to-peak amplitude of approximately 50 μV in at least 5 of 10 consecutive trials. We identified the muscle with the highest RMT and adjusted the stimulation intensity for the main experiment to 115 % of that muscle’s RMT. This was intended to reliably elicit MEPs while reducing the risk of encountering ceiling effects with MI related MEP modulation.

### TMS neurofeedback

Stimuli were created and presented using Psychophysics Toolbox-3 (Brainard, 1997) for Matlab (MathWorks, version 2013a) and additional custom scripts written in Matlab. Participants were instructed to keep their eyes open throughout the experiment and refrain from any muscle tensing or clenching. A trial consisted of a preparatory rest period, MI period, and a feedback period. Each trial started with a relaxation period to ensure that subsequently collected MEP amplitudes would not be influenced by changes in background muscle activation. During the relaxation period the root mean square of the EMG signal (rmsEMG) for each of the four muscles was tracked, within a sliding 500 ms window, and displayed to the participant in the form of four circles (Figure 1A). As long as the rmsEMG of all muscles remained below 7 μV, the circles would remain green and the trial would progress to its next stage. If, on the other hand, the rmsEMG of a particular muscle exceeded 7 μV, then the corresponding circle would turn red until rmsEMG dropped below 7 μV. The trial would proceed if the rmsEMG of all muscles was below 7 μV for 500 ms.

Once the rmsEMG criterion was met, the coloured circles were replaced by an instruction for which MI task had to be performed. The label “Rest” instructed the participant to rest and not to engage in any MI. Rest trials were used as baseline for providing feedback on the following MI trials. The label “Thumb”, “Index”, or “Pinkie” signalled the start of a MI task. Participants were instructed to kinaesthetically imagine movements with the signalled finger (target), without involving any of the other fingers (non-target). It is important to note that rmsEMG continued to be monitored during the MI period in the same way as in the preparatory rest period. A trial was automatically paused if the rmsEMG of any muscle exceeded 7 μV. The trial then only continued after the rmsEMG was below 7 μV for at least 500ms. If the rmsEMG remained sufficiently low throughout the trial, the MI phase lasted for a jittered interval of 4-6 seconds before a TMS pulse was discharged. This interval was jittered to prevent participants anticipating the pulse which is known to influence MEP amplitudes (Tran et al., 2021).

The feedback was provided 500 ms after the TMS pulse by displaying the normalized MEP amplitude for each right hand muscle. The normalized MEP amplitude of a given MI trial was obtained by dividing the MEP peak to peak amplitude of that trial by the averaged peak-to-peak amplitude across the preceding 10 rest trials. All baseline normalisations were done separately within the particular muscle: i.e. the FDI, APB, and ADM MEPs would be compared to the average FDI, APB, and ADM MEPs from the 10 preceding rest trials, respectively.

Normalized MEPs were shown by three bars representing the thumb, index finger and little finger MEPs, respectively. A horizontal white line represented for each muscle the average MEP amplitude across the 10 preceding rest trials. If the bar exceeded the white line, MEP amplitude during MI was larger than the resting MEP amplitude indicating facilitation. If the bar was below the white line, MEP amplitude during MI was below the resting MEP amplitude indicating inhibition.

Participants were instructed to adapt their MI strategy such that the target muscle bar would be strongly raised above the white line while the non-target muscle bars should be slightly below the white line (i.e. selective upregulating corticomotor excitability of the target finger muscle while downregulating the excitability of the two non-target finger muscles). As long as neither of the normalised non-target muscles MEP rose above 130 % of their respective baseline, and the target muscle MEP exceeded its baseline, a trial was deemed as successful and the participant received positive feedback, i.e. the bars turned green for 4 s. If the trial was unsuccessful the bars turned red.

### Control group

The control group underwent a similar protocol as the experimental group, except that the controls received non-informative neurofeedback. All feedback bars were of the same height and remained unchanged throughout the experiment. The proportion of positive and negative (green/red bars) feedback a control participant received, was matched to an individual of the experimental group. Such an approach ensured that visual feedback was matched across groups, while its informational content was different.

### Offline EMG data processing and analysis

Analysis was done using in-house scripts written in Python 3.6.7 (www.python.org) and R 3.6.3 (www.r-project.org) software.

Filtering was performed separately for the background EMG (bcgEMG), i.e. the interval before TMS was applied and MEP analysis in order to avoid “smearing” of the TMS artifact into the bcgEMG data. The bcgEMG was band-pass filtered between 30-800 Hz, with an additional notch filter at 50 Hz. Since the MEP amplitude is known to be affected by the bcgEMG (Devanne et al., 1997), we calculated the root mean square (RMS) of the bcgEMG within the interval of 105-5ms prior to the TMS pulse and excluded all trials where the bcgEMG RMS exceeded 7 μV. After applying this criterion, 3 % of the data needed to be discarded from further analysis. Furthermore, we used the RMS bcgEMG data for several control analyses: First, trials were pooled into two groups (target muscle vs. non-target muscles) and root mean square (RMS) of the bcgEMG was compared across sessions and between groups. Secondly, a Spearman correlation analysis was performed between normalised RMS of the bcgEMG and the normalised MEP data, within each session for each participant. BcgEMG was normalised in the same way as MEPs i.e. divided by the mean RMS bcgEMG activity of the preceding 10 rest trials. Thirdly, normalised bcgEMG was added as an additional regressor in all the MEP related mixed effects analysis, controlling thus for any potential influence BcgEMG might have had on the MEP modulations.

MEP onsets and offsets were detected using a BioSPPy library (Carreiras et al., 2015) which utilizes a thresholding procedure of 120 % of the mean signal + 2 standard deviations, in a sliding window starting from the TMS pulse. The following features were calculated from MEP time windows: normalised peak to peak (p2p) amplitude and the normalised RMS of MEP. MEPs with a peak-to-peak amplitude exceeding Q3 + 1.5 x (Q3 - Q1) were removed from further analysis, with Q1 denoting the first quartile and Q3 the third quartile computed over the whole set of trials for each participant. According to this criterion, 7% of the MEPs were removed from further analyses. Trials were pooled into two groups, target muscle vs. non-target muscles, and baseline normalised MEPs were compared across conditions, blocks, and between experimental groups. We compared performance using a metric called “Target Ratio”. The Target Ratio was calculated as a ratio of the normalised target muscle MEP to the normalised non-target muscle with the largest MEP 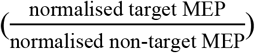. A Target Ratio larger than 1 implied selective modulation of corticomotor excitability.

### EEG acquisition

The EEG signal was recorded using 64 Ag/AgCl actiCAP active surface electrodes (Brain Products, Munich, Germany), an actiCHamp Plus amplifier and the Brain Vision Recorder proprietary software (Brain Products, Munich, Germany). The MNE Python library (Gramfort et al., 2013) was used for analysis. Electrodes were placed according to the extended international 10–20 system. All data were recorded at 1 kHz sampling frequency and all impedances were kept below 10–15 kΩ during the experiment.

### EEG pre-processing

We preprocessed the EEG data using the following steps: Data were first epoched from 3 s preceding the MI period up to 4 s of the MI period to remove the substantial artefacts that arise due to the TMS pulses. Subsequently, the data were down-sampled to 500 Hz, low-pass filtered at 100 Hz, and high-pass filtered at 0.1 Hz. We then performed Independent Component Analysis (ICA) using the ‘fastica’ algorithm, and removed components related to eye blinks and other artefacts. Epochs containing atypical noise component (e.g. caused by muscle activity) were rejected after visual inspection. The data were then re-referenced offline to an average reference. The epochs of interest were extracted from 1.5 s preceding the MI period up to 3.5 s of the MI period.

The time-frequency representation (TFR) was computed using the multitaper method. This method calculates spectral density for orthogonal tapers and averages them together for every channel/epoch. For further description of the general method see Percival and Walden (1993) and Slepian, D. (1978). The obtained power calculations were subsequently used in the event-related desynchronization (ERD) and representational similarity analysis (RSA) analysis.

### Event-related desynchronization analysis (ERD)

Event-related desynchronization (ERD) reflects a decrease of oscillatory activity related to MI processing. The trial epochs were baseline normalised by the mean activity of 1 s interval preceding the MI period. Given the importance of mu and beta band activity in MI related studies (McFarland et al., 2000) we examined these frequency bands in further detail and averaged power separately within the Mu (8 – 12 Hz), low beta (12-20 Hz) and high beta (20 – 28 Hz) bands.

Topoplots showcase relative power activity averaged either over the entire 3.5 s MI period (Figure 3A) or a more focused MI period of the last 1.5 s prior to the TMS pulse (Figure 4B), represented across 61 scalp electrodes.

### Representational similarity analysis (RSA)

Representational similarity analyses (RSAs; Kriegeskorte, Mur, and Bandettini 2008) were performed for both MEP and EEG measurements.

For the MEP based analysis, we used normalised peak to peak MEP amplitude and normalised RMS of the MEPs, extracted from each of the 3 right hand EMG electrodes. The normalisation was performed with respect to rest trials, as in the previous MEP analysis, and the resulting values were z-transformed over all trails. This resulted in feature vectors containing 3 electrodes (APB, FDI, ADM) x 2 parameters estimates (peak to peak, RMS) per trial which reflected the difference between MI and the preceding rest condition. We then used the feature vectors to calculate the cross-validated Euclidian distances between conditions. This was possible since every session included 3 blocks from which an equal amount of trials was randomly picked to form the crossvalidation folds. This process resulted in one representational dissimilarity matrix (RDMs) representing the inter-finger distances between conditions for each session and participant. From the RDM we then estimated the strength of the condition separation or “finger separability” by averaging the off-diagonal elements, capturing the distance between the MEP features for the three MI conditions by a single number. The more distinct MEP patterns were produced via MI, the higher the finger separability score.

An analogous RSA was performed for the EEG data. The preprocessed and power-transformed EEG data were subdivided into 3 power bands (mu, low and high beta bands). The power-transformed data were then averaged within each individual frequency band, resulting in one power value per electrode and per time point. The following process was repeated for each of the bands separately. Data were first z-transformed over all trials. The full channel dataset was then reduced to 15 electrodes which were defined a-priori and roughly covered sensorimotor-related areas in both hemispheres (‘CPz’, ‘C1’, ‘FCz’, ‘C2’, ‘C6’, ‘C4’, ‘Cz’, ‘FC3’, ‘P4’, ‘CP3’, ‘C3’, ‘FC4’, ‘C5’, ‘P3’,‘CP4’). The z-transformed data were then binned for each electrode over a 1 s window that was slid in 200 ms steps over the 5 s period, i.e. the last 1.5s of the preparatory rest period and the first 3.5s of the MI period. This resulted in 25 overlapping time windows each comprising of averaged data within 1 s time bin. To obtain the condition-specific activations for each bin and electrode, we constructed a design matrix using dummy coding for each task condition (thumb, index, pinkie) and estimated condition-specific beta coefficients by means of a linear regression model for each electrode (Luyckx et al., 2019). This revealed 3 beta coefficients for every electrode and every time bin. We then calculated the cross-validated Euclidian distances between conditions. An equal amount of trials were randomly picked from the three blocks per session to form the crossvalidation folds. This process resulted in 25 representational dissimilarity matrices (RDMs) representing the inter-finger distances between conditions throughout the 5 s interval. These calculations were repeated for every session and every participant. From each RDM we estimated the finger separability score by averaging across the off-diagonal elements.

Next, we explored the evolution of finger separability within a trial. To this end the RDMs were temporally smoothed via linear interpolation and the finger separability progression was displayed throughout the duration of 5 s, covering the preparatory rest period (1.5s) and the MI period (3.5s). In order to visualize inter-finger distances in a 2-dimensional (2D) plane we applied multidimensional scaling (MDS; Buja et al. 2008). Multidimensional scaling is a dimensionality reduction algorithm that projects values into a smaller dimensional space while aiming to preserve the relative distances. To remove the arbitrary rotation induced by Multidimensional scaling, we performed Procrustes alignment. The Generalized Procrustes analysis compares and aligns shapes obtained for different timepoints, sessions or participants to an averaged shape (Grice and Assad, 2009). This process left us with triangular shapes of neural patterns, where each edge displays the distance between two conditions. Using this representation, two MI conditions are more separable when they are further apart. We present multidimensional scaling plots of group-level distances for the last 1.5s of preparatory rest period and the last 1.5 s of MI period (i.e. 2s – 3.5 s; Figure 5).

For the final analysis, we estimated finger separability by averaging the RDMs for the 1.5 s of the preparatory rest period and the 2 – 3.5 s interval of the MI period. The preparatory rest finger separability scores were subtracted from the MI related finger separability scores, providing thus an index of how strongly finger separability was changed by MI. Statistics were calculated to compare these values between groups and sessions (Figure 6).

### Statistical procedures

Statistical inferences were drawn using Mixed Effects Models estimated using R (R Core Team, 2014) package glmmTMB (Brooks et al., 2017) as they offer greater flexibility for modelling repeated measures over time in comparison to traditional ANOVA methods (Gueorguieva and Krystal, 2004). Fixed effects were Group (NF vs Control) and either Sessions (1-5), or Blocks (blocks 1-15). Participants were modelled as random effects. Results are reported as fixed effects, Wald Chi-square (*X*^2^) test statistic and the corresponding p-values. The significance level alpha was set to 0.05 for all inferential tests (if not stated otherwise). In cases of multiple comparisons, p-values were corrected using the false discovery rate (FDR) with q set to 0.05.

Statistical analysis of spatiotemporal EEG data and time-frequency plot comparisons were performed using cluster-based analysis. Significant clusters were identified using cluster-corrected nonparametric permutation tests (iterations = 1000, cluster-level thresholds at p<0.05, unless stated otherwise) (Maris and Oostenveld, 2007).

## RESULTS

We investigated whether TMS-NF training promotes selective MEP modulation measured across the thumb, index and little finger muscles. Participants who underwent five sessions of successive TMS-NF training were compared to a control group which completed the identical protocol but received uninformative feedback.

### TMS-NF training promotes selective modulation of corticomotor excitability

We first compared how MEP amplitudes changed in the NF versus control group during the course of training. We found that the two groups differentially modulated their target and non-target MEP amplitudes: the NF group successfully adapted their MI such that MEPs of the target muscle where upregulated, while MEPs of the non-target muscle were downregulated. This selective facilitation of target finger muscles and inhibition of non-target finger muscles during MI became gradually larger from session 1 to 3 indicating learning across the blocked sessions. The difference between MEP amplitudes of target versus non-target muscles was maintained in the interleaved sessions 4 and 5, however, the difference was less pronounced. In the control group, there was a trend that MEPs were larger in the target than in the non-target muscles, however, this difference was much less pronounced than in the NF group and changed only slightly during the course of training. Accordingly, the block (1-15) x muscle (target, non-target) x group (NF, control) mixed effects analysis revealed a significant 3-way-interaction (X^2^(14,10)=25.6, p=0.029).

Next, we repeated the statistical analysis separately for session 1-3 and session 4-5. We confirmed the 3-way interaction for session 1-3 (X^2^(8,10)=18.19, p=0.019) but not for session 4-5 (X^2^(5,10)=1.72, p=0.881) which only revealed a significant muscle x group interaction (X^2^(1,10)=10.01, p=0.001). This implies that the largest learning gains regarding MEP modulation were achieved across session 1-3 indicating that participants acquired the ability to selectively facilitate single finger representations using MI. Crucially, participants were still able to successfully modulate MEP amplitudes during the interleaved sessions when compared to controls. One unexpected observation was that the NF group performed better than the control group already during the very first training block. A more fine-grained analysis of the first block (figure 2B) shows that MEP modulation was relatively similar in both groups for the first 21 trials. After that, the NF group became much more successful with facilitating the target muscle, a phenomenon which was not observed in the control group. This suggests that the TMS-NF was highly intuitive for our participants and could be quickly employed to adjust MI strategies.

**Figure 2.**
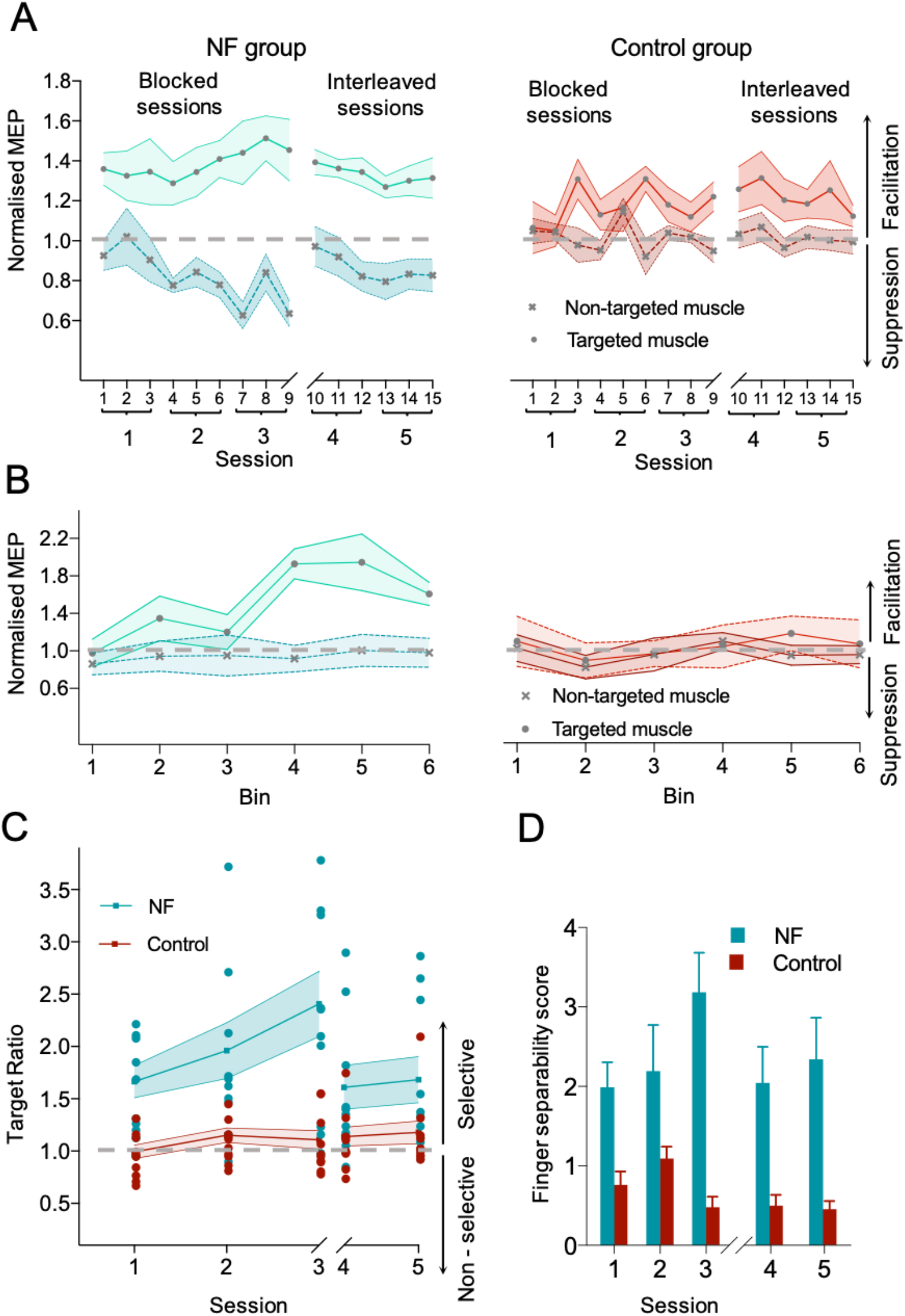
TMS-NF promotes finger selective modulation of corticomotor excitability. A) Selective MEP modulation was bidirectional, i.e. participants that underwent TMS-NF training (left panel) learned to upregulate the excitability of the target muscle (filled circles) while simultaneously downregulating the excitability of non-target muscles (cross). while the controls (right panel) did not show such pattern. B) Detailed analysis of the first block of session 1. The 42 trials in block 1 were binned into groups of 7 trials, resulting in 6 bins. Neither the NF groups (left panel) nor the control group showed selective facilitation of the target muscle for bins 1-3. However, for the NF group only, the target muscle was substantially upregulated from bin 4 onwards. C) Changes of the target ratio in the NF group (red) and controls. Values exceeding a target ratio = 1 indicate muscle-selective facilitation. Circles show individual subject data. Learning, operationalised as a session x group interaction effect (p=0.017), took place over sessions 1-3. The NF group did not improve further in sessions 4-5, but still maintained superior performance in selective modulation in these sessions compared to the control group (group effect; p=0.019). Note that the task in sessions 4-5 was more challenging as the trials were organised in an interleaved manner (i.e. any type of MI could appear within a block). D) MEP-based representational dissimilarity analysis. Separability scores are shown in arbitrary units which higher scores indicating higher separability based on features extracted from the TMS evoked responses. Finger separability was significantly higher in the NF compared to the control group (p<0.001) and this difference changed with training (group by session effect; p=0.017), indicating TMS-NF promotes increasingly dissociable excitability patterns underlying single finger representations. This result is in accordance with our Target Ratio results that show reduced TR in the interleaved sessions, probably reflecting increased task difficulty.

**Figure 3.**
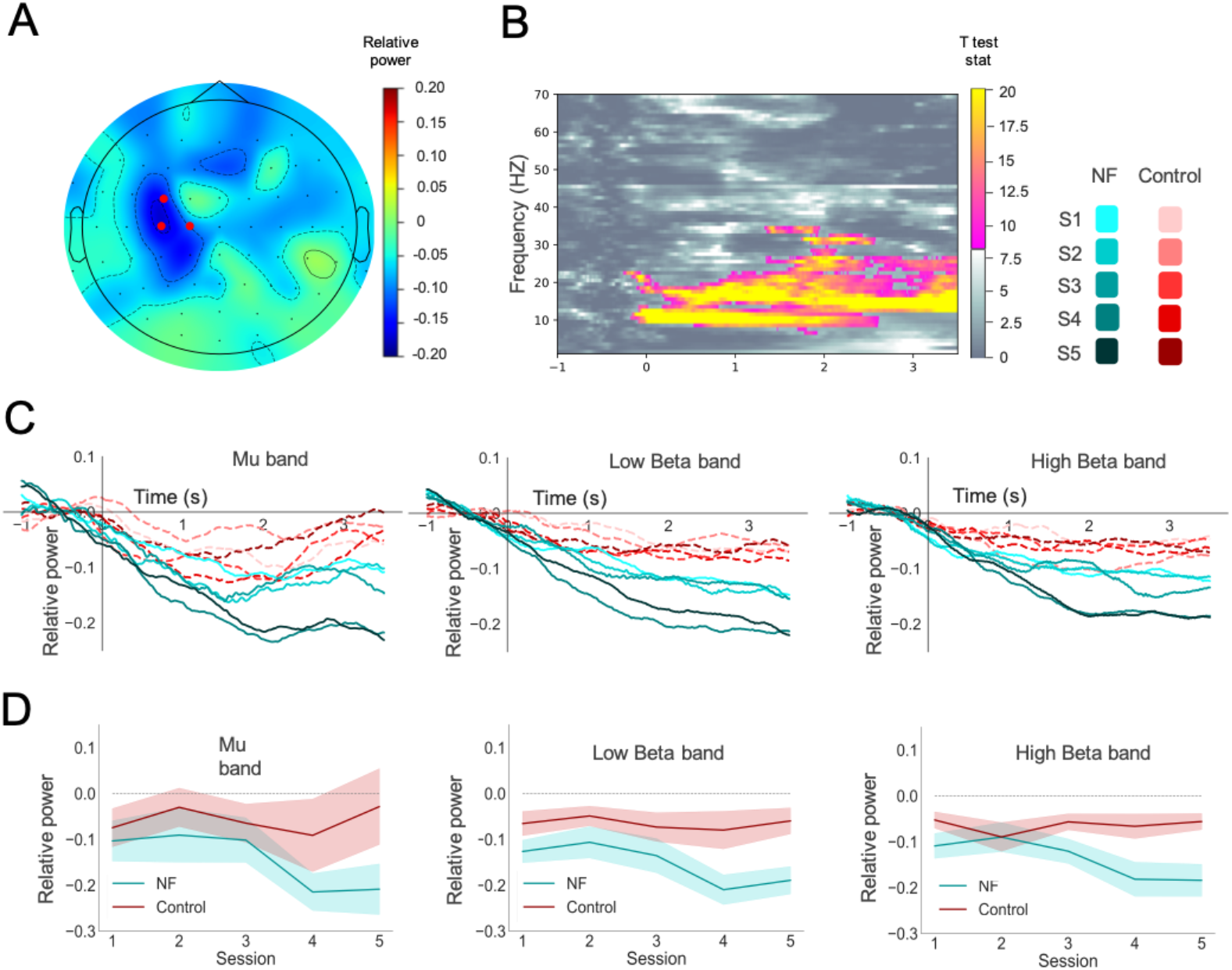
ERD over sensorimotor areas. A) The topoplot shows changes in relative power from the preparatory rest period to the first 3.5s of the MI trials within a broad frequency band containing several sensorimotor rhythms (8-28 Hz). Data were collapsed across groups and conditions. Darker blue colors indicate larger event related desynchronization (ERD) based on relative power. Red dots indicate a cluster of electrodes which shows significant ERD relative to baseline (p<0.001). B) Time-Frequency analysis was performed on the mean activity from the cluster identified in panel A. Bright colored sections illustrates frequencies that exhibit significant ERD (p<0.001) while the gray colors indicate the non-significant sections. C) Line plots illustrate changes in relative power across Mu (8-12 Hz), Low (12-20 Hz) and High beta (20-28 Hz) bands in all 5 sessions for neurofeedback (teal) and control (red) group. There is a consistent ERD pattern present across majority of sessions. Strongest ERD is observed in the period of 2s – 3.5 s after MI has been cued (corresponding to 0s). D) Mean activity from electrodes comprising the cluster identified in A) was plotted across 5 sessions in mu, low beta and high beta frequency band (extracted from 2-3.5 seconds after MI has started). Shaded areas indicated standard errors of the mean. A significant Group effect was observed (p_FDR_<0.001) for all three bands implying larger ERD in the NF group compared to controls.

**Figure 4.**
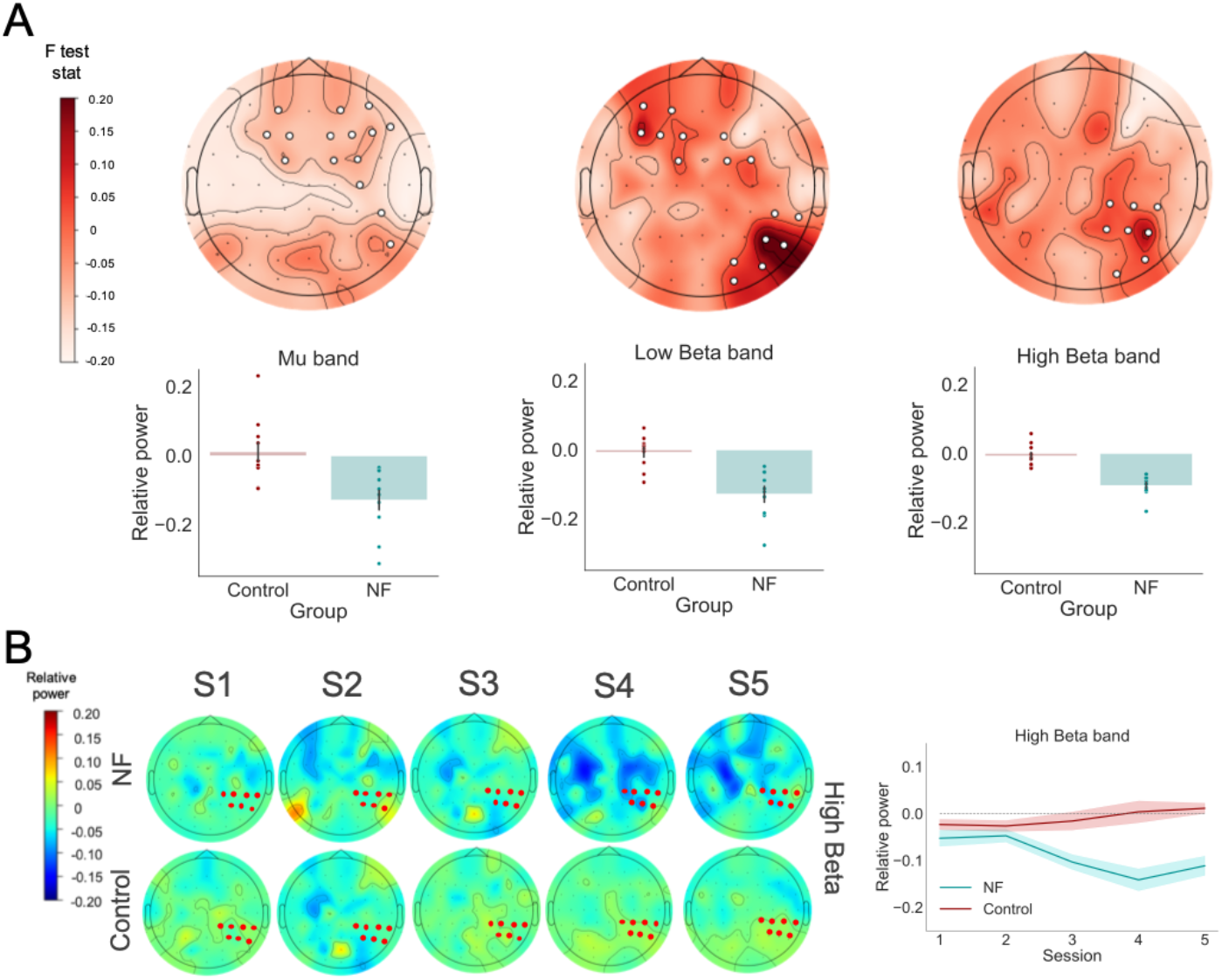
Brain wide analysis of group and session effects for sensorimotor rhythms. Desynchronisation was quantified between the rest period preceding each MI trial and the period from 2-3.5s after the MI condition has been cued. A) Cluster-based main effect of group. White dots indicate electrode clusters which exhibit stronger desycnhronisation in the NF compared to the control group in the mu band (p_cluster_=0.1), low beta band (p_cluster_=0.008, p_cluster_=0.013) and high beta band (p_cluster_=0.008) that showed difference between the groups across five sessions. The bar plots indicate relative power differences between groups averaged over all electrodes in a particular cluster. Dots indicate individual participants. B) Cluster-based group x session interaction effect in the high beta band. Topoplots depict changes in the relative power for the NF (upper row) and control group (lower row) across 5 sessions. In the NF group, ERD becomes progressively stronger and more distributed with additional sessions. This is not the case in the control group. Marked electrodes signify an electrode cluster in the high beta frequency band that showed a significant group x session interaction effect. The right panel shows changes in high beta band synchronisation averaged across all cluster electrodes and plotted for each group over all sessions with the shaded area illustrating standard errors. No interaction effects reached significance for the two other bands.

**Figure 5.**
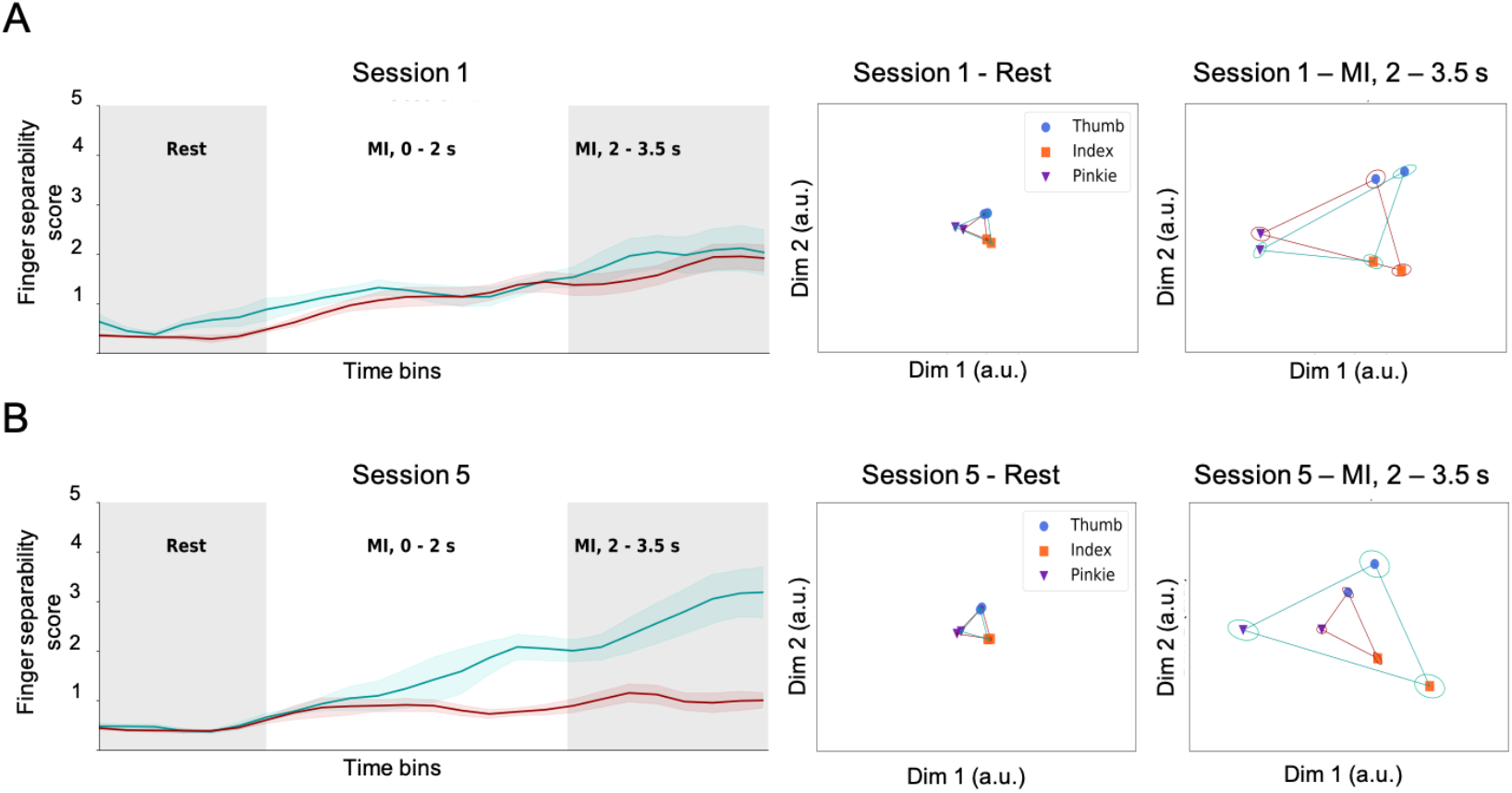
Evolution of EEG-based finger separability. A) The left panel illustrates the evolution of finger separability within a trial in session 1 averaged within the NF group (blue) and controls (red). Finger separability scores (see methods for details) are expressed as a distance in arbitrary units. As expected, distance scores were rather small during the preparatory rest period and gradually increased during the MI period. In session 1 there were no significant differences between the groups. The mid and right panel illustrates inter-finger difference patterns projected into the same 2D space. Larger distances between the finger representations indicate larger separability of the finger specific EEG patterns. Ellipses indicate standard errors. The mid panel shows that EEG patterns were similar during the preparatory rest period (distances between the different finger representations is small) but became more distinct during the 2 – 3.5s interval of the MI period. B) The evolution of finger separability within an averaged trial in session 5. The two groups did not differ during the preparatory rest period, but as MI unfolded, finger separability was substantially greater in the experimental group than in the control group. The increase in inter-finger distances was similar across all 3 finger MI representations.

**Figure 6.**
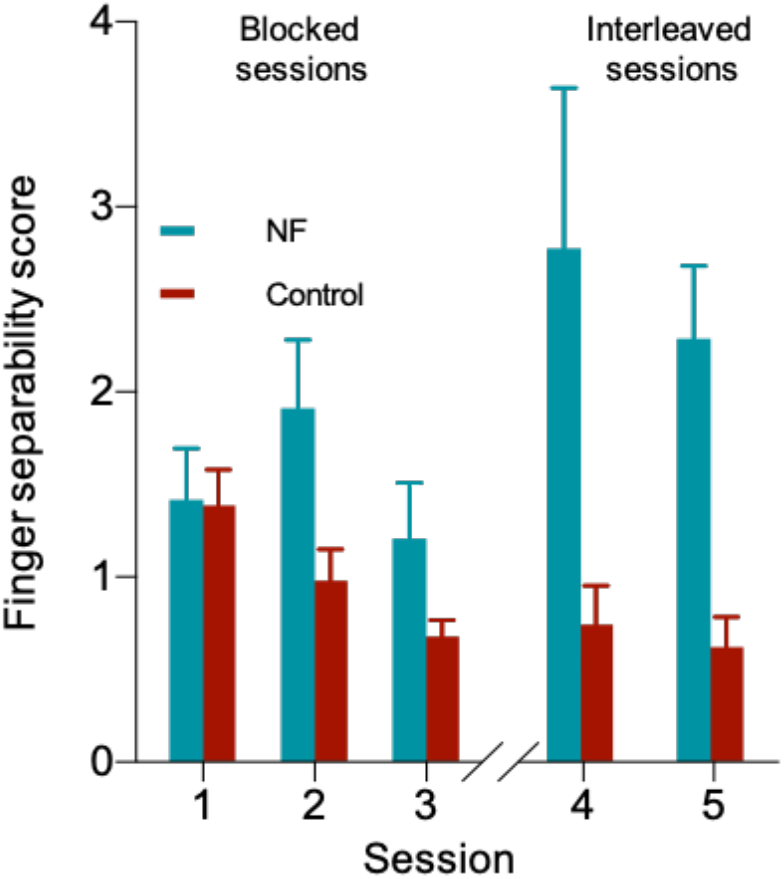
Separability of neural patterns underlying single finger representations. An EEG-based representational dissimilarity matrix (RDM) was calculated to extract inter-finger distances. We calculated a finger separability score by averaging all unique off-diagonal values in the inter-finger RDM and compared this score between groups and across the sessions. While the finger separability predominantly increased across sessions in the NF group, the Control group displayed an opposite pattern. Sessions 4-5 revealed the greatest difference in finger separability between the groups.

We expressed the above MEP modulation by the Target Ratio (TR) with TR > 1 indicating a high level of selectivity in modulating the MEPs of target versus non-target muscles. The NF group had a higher Target Ratio (average TR_NF_ = 1.86) compared to the control group (average TR_control_ = 1.11), confirmed by a significant group effect (X^2^(1,10)=34.96, p<0.001), and this between-group difference increased across sessions, confirmed by a significant interaction effect (group x session effect, X^2^(4,10)=13.81, p=0.007). Additionally, mixed effects revealed that this difference in TR between groups increased across sessions 1-3 (group x session effect, X^2^(2,10)=8.08, p=0.017) and remained stable over session 4-5 (Group effect X^2^(1,10)=5.47, p=0.019). Therefore we conclude that participants acquired the skill of selective corticomotor modulation using MI during session 1-3 while in session 4-5 they successfully displayed their selective modulation ability without further improvement relative to the control group.

Even though we excluded trials with high EMG activity, even small modulation of background EMG might influence MEP amplitudes (bcgEMG, Devanne et al., 1997). An additional control analysis revealed no evidence suggesting that the above results might have been driven by modulations in background EMG (Supplementary figure 2).

Finally, we were interested whether TMS-NF training leads to increased inter-finger separability of the MEP patterns. Therefore, we determined finger separability scores based on RDMs, which were then compared across sessions and between groups. Note that, in contrast to the Target Ratio analysis, RDMs do not make a-priory assumptions on how the MEP pattern would relate to the three MI conditions. We found that finger separability based on the MEP pattern was higher in the NF than in the control group (Figure 2D). This difference was already present in the first session but, importantly, it further increased with additional TMS-NF training (Group x Session, X^2^(4,10)=12.01, p=0.017).

### Event related desynchronization is modulated by TMS-NF training

We recorded EEG activity during all sessions and examined how MI-related brain activity changed during the course of TMS-NF training. Therefore we determined the event-related desynchronisation (ERD) of neural activity in the mu, low and high beta band, which is a well-known and robust EEG-based marker of neural processes related to MI (McFarland et al. 2000).

According to our a-priori hypothesis, we first investigated whether ERD in sensorimotor areas would differ between the NF and control group. Therefore, we tested which electrodes exhibited significant activity changes during a MI trial irrespective of the MI finger condition, session or group effects. We analysed which electrodes would exhibit a significant ERD effect in the broad 8-28 Hz frequency spectrum during the first 3.5s of the MI period. Note that this period was not confounded by the TMS artifact since the TMS pulse was delivered not earlier than 4 s after MI has started. A cluster-based permutation test (p_cluster_<0.001) revealed pronounced ERD for the electrodes C1, C3, FC3 which are likely to reflect activity in left sensorimotor cortex. Next, we averaged the activity of those electrodes and created a time frequency plot covering 4-70 Hz for 1 s prior to the start of MI and the first 3.5 s of MI. This wider frequency range was chosen to explore whether MI might influence other frequency bands than the hypothesized SMRs. Clusters of significant ERD effects (p_cluster_<0.001) were mainly detected in the mu (8-12 Hz) and beta (12-28 Hz) bands (Fig 3 B) covering the whole 3.5s MI period which was analysed. All subsequent analyses focused on the mu (8-12 Hz), low beta (12-20 Hz) and high beta bands (20 – 28 Hz) i.e. the cortical rhythms which have most frequently been associated with MI (Kim et al., 2016; Shahid et al., 2010; Tariq et al., 2020). We further examined how the ERD of the NF and the control group evolved during a trial and across sessions (Figure 3C). We found that ERD increased gradually over time. Particularly for the beta bands, the strongest ERD effects were observed at 2 - 3.5s after MI was initiated by the participants. Moreover, ERD of the NF group became gradually stronger across the sessions 1-5 with the largest ERD emerging during sessions 4-5 (Figure 3C, teal line plots). In the control group, by contrast, no such progression across sessions was visible (Figure 3C, red line plots).

To explicitly test the evolution of ERD in further detail, we averaged the ERD data within the 2 - 3.5s interval and compared these values between groups and sessions (Figure 3D). Separate analyses for the mu, low beta and high beta bands revealed a Group effect (mu, X^2^(1,10)=10.77, p=0.001; low beta, X^2^(1,10)=24.54, p<0.001; high beta, X^2^(1,10)=26.41, p<0.001) confirming that ERD was stronger in the NF than in the control group. Even though the figure shows a trend that ERD is strongest in session 4-5 for the NF but not for the control group, the Group x Session interaction effect did not reach significance, potentially because clear group differences are already present in session 1 (mu, X^2^(1,10)=2.94, p=0.57; low beta, X^2^(1,10)=2.92, p=0.56; high beta, X^2^(1,10)=5.73, p=0.22). In summary, single finger MI caused ERD of the mu and beta rhythms in electrodes overlaying sensorimotor areas. ERD was significantly stronger in the NF than in the control group.

Next we asked whether other electrodes might reveal Group or Group x Session effects regarding MI induced ERD in the three frequency bands of interest. Therefore, we employed separate permutation-based spatiotemporal cluster analyses for the mu, low beta and high beta band over all electrodes, testing either for a group main effect or session by group interaction (Figure 4). A trend towards higher mu ERD in the NF than the control group was detected for a bilateral cluster containing mainly fronto-central electrodes (p_cluster_=0.100; Figure 4A). For the low beta band, a significantly higher ERD was detected over bilateral fronto-central cluster (Group effect, p_cluster_=0.013) which party overlapped with the electrode cluster detected for mu ERD. Additionally, there was a significant group difference in low beta band ERD for a right parieto-occipital electrode cluster (Group effect, p_cluster_=0.008). For both clusters ERD was stronger in the NF than the control group (Figure 4A). Finally, for the high beta band, we detected a highly similar right parieto-occipital cluster as for the low beta bend, where ERD was significantly stronger in the NF than the control group (p_cluster_=0.008; Figure 4A).

Interestingly, similar clusters were detected to exhibit a trend towards a group x session interactions effect (Supplementary figure 3) indicating that ERD became gradually stronger in the NF group while remaining unchanged in the control group. However, only one cluster containing predominantly parietal electrodes of the right hemisphere survived correction for multiple comparisons when tested for high beta band activity (p_cluster_=0.014; Figure 4B). Thus, even though TMS-NF training caused an increase of ERD in electrodes overlaying left sensorimotor cortex, a fronto-central cluster and a right parieto-occipital electrode cluster also exhibited strong group differences. Particularly for the parieto-occipital electrode cluster, high beta band ERD increased progressively with training but only for the NF group. These results suggest that TMS-NF might have influenced activity in higher-order areas involved in sensorimotor processing.

### TMS-NF training leads to increased inter-finger separability

While the average ERD changes reveal an overall effect of MI on the related neural activity between groups, they provide little information about the potential differences between the MI conditions and how they evolve with TMS-NF training. Since we were interested in how TMS-NF training may improve the separability of neural patterns of individual finger MI, we employed representational similarity analysis (RSA; Kriegeskorte et al.,2008). RSA is a multivariate method that can be used to infer whether different MI tasks are represented by specific neural activity patters. To do so, specific features are extracted from the neural activation patterns caused by MI of different finger movements and represented within an abstract feature space. The Euclidean distance metric is used to quantify the finger separability i.e. whether these representations are similar (i.e. located close together in the feature space) or different (i.e. more distant in feature space).

First, we were interested in the evolution of inter-finger differences within a trial. We found that the finger separability scores were rather small during the preparatory rest period but gradually increased once MI has started, indicating that the EEG activation pattern becomes more and more separable when thumb versus index versus little finger MI is performed (Figure 5). Importantly, finger separability showed no differences between the groups in the rest period (X^2^(1,10)=1.48, p=0.22) and increased towards the end of MI period (main effect of phase i.e. from preparatory rest to MI; X^2^(1,10)=108.99, p<0.001). For session 1, this increase of separability during the MI period was similar in both groups (Figure 5A). In session 5, however, separability during MI was clearly more pronounced in the NF group than in the control group (Figure 5B).

Furthermore, we compared finger separability scores across sessions and between the two groups. To account for any bias that could influence our conclusions, but stems from inter-finger distances already present at rest, we subtracted out the preparatory rest period finger separability and compared the remaining average finger separability score between 2-3.5s across sessions and between groups (Figure 6). The groups exhibited similar finger separability caused by MI in session 1 (p_FDR_=0.90) but, crucially, the finger separability increased significantly more across sessions in the NF group when compared to the control group (Figure 6; Group x session interaction X^2^(4,10)=17.55, p=0.001). In fact, RDMs decreased slightly in the control group such that the group difference was strongest in sessions 4 and 5 (session 4 p_FDR_=0.01; session 5 p_FDR_=0.001). Our finding implies that single finger MI initially activated largely similar neural activity patterns (session 1), but as participants underwent repeated TMS-NF based training aimed to tune their MI strategies, these EEG patterns became more separable, suggesting that a higher level of finger individuation was reached during MI.

## DISCUSSION

We aimed to investigate whether participants can be trained to selectively facilitate single finger representations via motor imagery when neurofeedback is provided on the basis of TMS-evoked motor potentials. We show that TMS-NF training enables participants to upregulate corticomotor excitability of the target finger while simultaneously down-regulating corticomotor excitability of non-target fingers. This ability for “finger individuation” during MI was associated with strong desynchronisation of sensorimotor brain rhythms, most notably within the beta band, which was observed in a left sensorimotor cluster, a bilateral-frontal electrode cluster and a right parieto-occipital electrode cluster. Additionally, neural activation patterns underlying single finger MI were shown to be more dissociable when participants were trained with TMS-NF than when no such feedback was provided.

### TMS-NF promotes selective modulation of corticmotor excitability

In general, NF is an operant conditioning-based technique in which individuals are reinforced to modulate their mental states. Numerous studies illustrate successful application of NF in the neuroscience domain. Its effectiveness is demonstrated on micro- (electrocorticographic; Gharabaghi et al. 2014), meso- (fMRI; Scharnowski et al. 2012) and macro-scale (electrocorticographic; Omejc et al. 2019) levels, both in healthy and clinical (Wyckoff and Birbaumer, 2014) settings.

Here we used TMS-NF which is still a relatively new technique. Nonetheless first studies showed that it can be instrumental in obtaining voluntary control over the excitability of a motor system, e.g. by facilitating (Koganemaru et al. 2018; Ruddy et al. 2018) or inhibiting (Majid, Lewis, and Aron 2015) MEPs over a wide range of values. Our study extends these previous results by showing that TMS-NF can help participants to selectively activate individual finger representations via MI. More specifically, TMS-NF training allowed participants to volitionally facilitate cortical representations of the target finger (by approx. 40%) while simultaneously suppressing corticomotor excitability of the other fingers (by approx. 20%). It is important to note that these results were not driven by changes in background EMG activity, which was closely controlled throughout the experiment and further inspected via several control analysis.

Additionally we found that selective TMS-NF was quickly utilized by the participants as indicated by more than 50 % difference in Target Ratio between the two groups already in the first session. This unexpected result suggests that TMS-evoked neuroelectrical signals offer an opportunity for providing intuitive and robust feedback even if selective MI strategies need to be developed. This might be advantageous, particularly when participants have to perform complex MI task which typically require multiple training sessions (Rogala et al., 2016). Moreover, one has to keep in mind that during the first three sessions MI conditions were blocked, i.e. the participants could re-activate the same MI state several times in a row. In session 4 and 5, by contrast, the required finger-specific facilitation/suppression pattern varied on a trial-by-trial basis. We observed a clear drop of performance in the NF group, as assessed by the Target Ratio, indicating that constantly switching between MI conditions imposes high demands. Based on our data, we cannot predict whether additional sessions would result in an increase in selective corticomotor modulation proficiency, on this more difficult level, that would eventually match the blocked sessions success. Furthermore, not all participants were able to selectively modulate their corticomotor excitability. One participant did not improve performance with training (average TR= 0.95). This could be attributed to the known “illiteracy” phenomenon reported in brain computer interface (BCI) studies, where as much as 15-30 % of participants are found to be non-responders to BCI training (Blankertz et al., 2010). Moreover, it has been shown that strong corticomotor excitability and selective MEP modulation is linked to an individual’s ability to perform vivid Motor Imagery as assessed via a combination of psychological, behavioural and psychophysiological measurements (Lebon et al., 2012). Accordingly, it is possible that TMS-NF is not effective in individuals who have difficulties with vivid motor imagery per se.

Selective modulation of single finger representations by MI, or “mental finger individuation”, may have implications for stroke rehabilitation and recovery. The ability to move individual fingers while keeping the non-moving ones stationary (Schieber, 1991) is a complex motor task which requires a high level of cortical control. Accordingly, the ability for finger individuation is impaired when the corticomotor tract or cortical motor areas are damaged. For example, in stroke patients it has been shown that finger individuation measurements are highly correlated with both Fugl-Meyer Assessment (FMA; Fugl-Meyer, 1975), and the Action Research Arm Test (ARAT; Lyden and Lau 1991). While the feasibility of TMS-NF training after stroke has already been demonstrated in a clinical setting (Liang et al., 2020), it remains to be tested whether TMS-NF training targeting the selective modulation of hand or finger representations via MI could aid recovery.

### Mental finger individuation is related to increased desynchronisation of mu and beta band activity

MI is known to be associated with changes in SMRs over the sensorimotor cortex contralateral to the imagined effector (Nam et al., 2011). Our results confirmed these previous findings but also provide new evidence that shaping MI by increasing corticomotor excitability for one finger representation while simultaneously suppressing other finger representations caused substantial desynchronisation in the mu and beta band. This desynchronisation was particularly strong for session 4 and 5, i.e. when a new target finger had to be selectively imagined on a trial-by-trial basis. This finding is in line with previous reports, indicating that particularly beta desynchronization becomes stronger when demands on movement selection increase during MI (Brinkman et al., 2014). While both rhythms desynchronized also in the control group, this effect was much smaller and did not improve across sessions, confirming that MI related cortical changes are typically minor when no reliable feedback is provided (Holper and Wolf, 2010). Several previous studies have shown that combining MI training with EEG neurofeedback can enable participants to volitionally control mu (Takahashi et al., 2018; Takemi et al., 2013) or beta (Kraus et al., 2016) band activity within sensorimotor areas. Moreover, it has been shown that when participants control their brain state such that one of these sensorimotor rhythms is strongly desynchronized, corticomotor excitability of the modulated primary motor cortex is high when probed with TMS (Kraus et al., 2016; Takahashi et al., 2018; Takemi et al., 2013). Our results extend these findings by showing that the relationship between SMR desynchronisation and corticomotor excitability is bidirectional, i.e. volitional control of the corticomotor excitability state is associated with strong desynchronisation of SMRs.

Although the majority of previous studies report desynchronisation of SMRs predominately contralateral to the imagined effector (Nam et al., 2011; Takemi et al., 2013), we detected desynchronisation of SMRs in mid-frontal electrodes and in a cluster of parieto-occipital electrodes in the hemisphere ipsilateral to the imagined effector. This indicates that MI training guided by TMS-NF not only affects SMR activity in electrodes overlaying contralateral sensorimotor areas but also other brain regions, potentially in secondary motor areas within frontal cortex, and higher order areas in the right hemisphere. Note, however, that we did not perform source localisation such that this anatomical interpretation is highly speculative and subject to future research.

### Selective TMS-NF based training leads to increased separability of neural patterns underlying MI

Our RSA results are illustrative of the potential influence TMS-NF training can have on the separability of neural patterns underlying single finger MI. Importantly, higher separability in the NF compared with the control group was observed when either MEP or EEG features extracted from electrodes overlying bilateral sensorimotor cortex were used. This indicates that TMS-NF reinforces mental strategies that would eventually result in more dissociable EEG patterns for finger-specific MI.

Visualisation of EEG related finger separability allowed us to observe how inter-finger distances gradually increased throughout the course of an average trial. Importantly, distances were small and completely indistinguishable between the groups in the preparatory rest period and became larger and more separable towards the end of MI period. This effect was only present in the advanced stage of MI (2 - 3.5 s), whereas the divergence was mostly absent in the initial part of the MI period (0 – 2 s). Separability developed differently across groups and sessions when analysing MEP-based versus EEG-based features: MEP derived finger separability was already in the first session higher in the NF than in the control group (p_FDR_<0.001). EEG related finger separability, by contrast, were similar between the two groups in the first session (p_FDR_=0.90), but diverged strongly in session 4 and 5 (session 4 p_FDR_=0.01; session 5 p_FDR_=0.001). This effect resulted not only from an increased separability across sessions in the NF group but partly also from decreased separability in the control group, which might have arisen from diminished motivation when faced with a repetitive activity deprived of informative feedback. It is possible that the EEG-based separability estimates were influenced by higher movement selection demands in session 4 and 5 where participants had to switch between the different finger representation on a trial-by-trial basis. It is interesting that TMS-NF not only evoked increasingly dissociable MEP patterns, but subsequently resulted also in more separable EEG-related neural patterns. These findings might prove relevant for future brain computer interface (BCI) designs. A great deal of current studies, aiming to perfect BCI related techniques, focus on optimising aspects of the decoding algorithms (for example, Al-Saegh, Dawwd, and Abdul-Jabbar 2020; Roy et al. 2019). Many sophisticated methods (Aggarwal and Chugh, 2019) have already been tried with limited success in surpassing a seemingly fixed plateau, determined by the inherently limiting signal to noise ratio of the EEG signal. On the other hand, approaches that try to instruct and incentivise the users to learn how to self-modulate their neural patterns towards increased separability, i.e. to improve signal to noise ratio, are greatly underrepresented. The evidence presented here suggests that TMS-NF training of fine finger MI could be a valuable new approach to further improve the performance of state-of-the-art BCI systems, however, therefore it needs to be directly compared to an EEG-based NF approach.

## Conclusion

In conclusion, we show that TMS-NF is a new approach which enables participants to learn how to mentally individuate their fingers by selectively modulating corticomotor excitability of intrinsic hand muscles. Changes in the ability to volitionally control finger specific corticomotor excitability were associated with increasing desynchronisation of sensorimotor rhythms recorded in electrodes overlaying contralateral sensorimotor cortex but also in a mid-frontal and an ipsilateral parieto-occipital electrode clusters. Interestingly, separability of EEG activity recorded over bilateral sensorimotor cortex was significantly higher in the NF group than in the control group during the last session where the mentally individuated finger had to be newly selected on a trial-by-trial basis. Our findings suggest that selective TMS-NF might constitute a new tool for facilitating functional recovery after stroke in addition to physical training.

## Acknowledgements

This work was supported by the Swiss National Science Foundation (320030_175616) and by the National Research Foundation, Prime Minister’s Office, Singapore under its Campus for Research Excellence and Technological Enterprise (CREATE) programme (FHT). We thank André Höpli and Yves Baumgartner for their assistance during data collection.

## Data availability statement

Data are openly available on the ETH Library Research Collection with the DOI: http://hdl.handle.net/20.500.11850/468729.

