## Supplementary Material for "Mental individuation of imagined finger movements can be achieved using TMS-based neurofeedback"

Supplementary figure 1

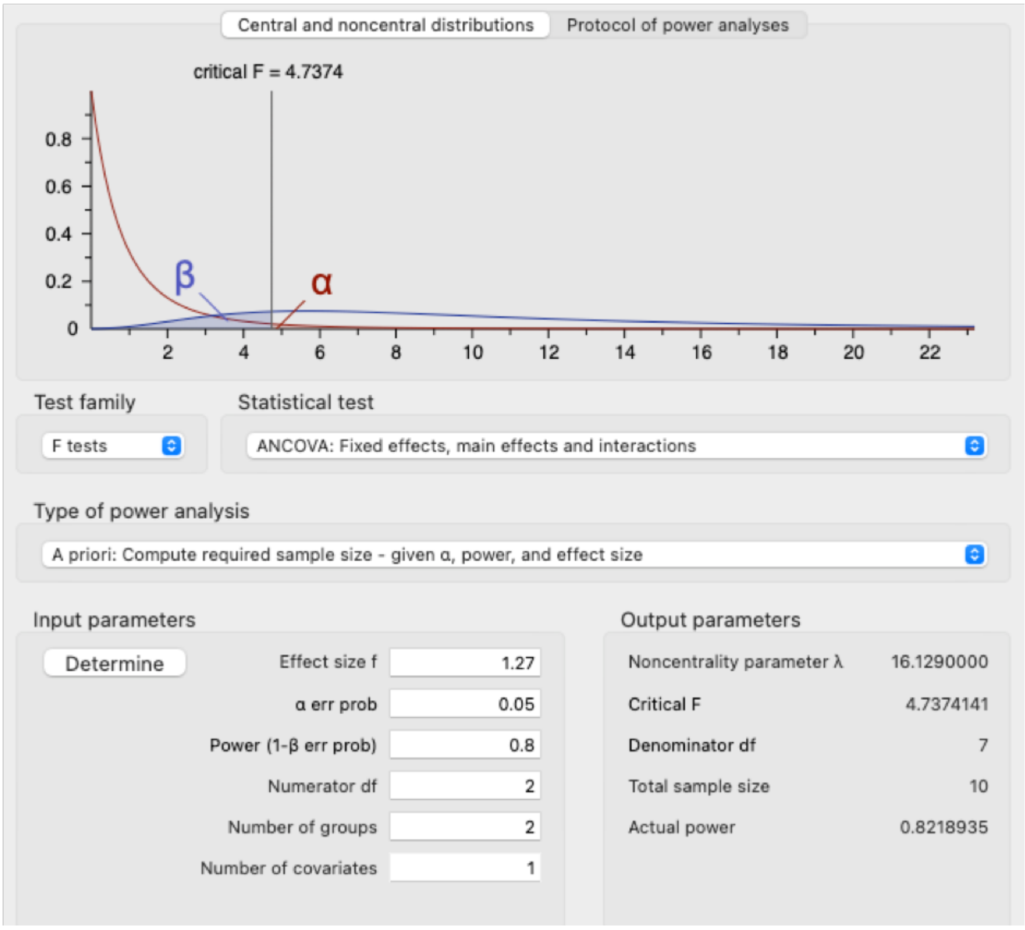


***Supplementary figure 1***. *We conducted sample size calculation using G*Power software version 3.1, based on the alpha value of 0.05, inclusion of two groups and an effect size of 1.27 (effect size reported in Ruddy et al., 2018) reported on the between-group differences in MEP modulation.*

Supplementary figure 2

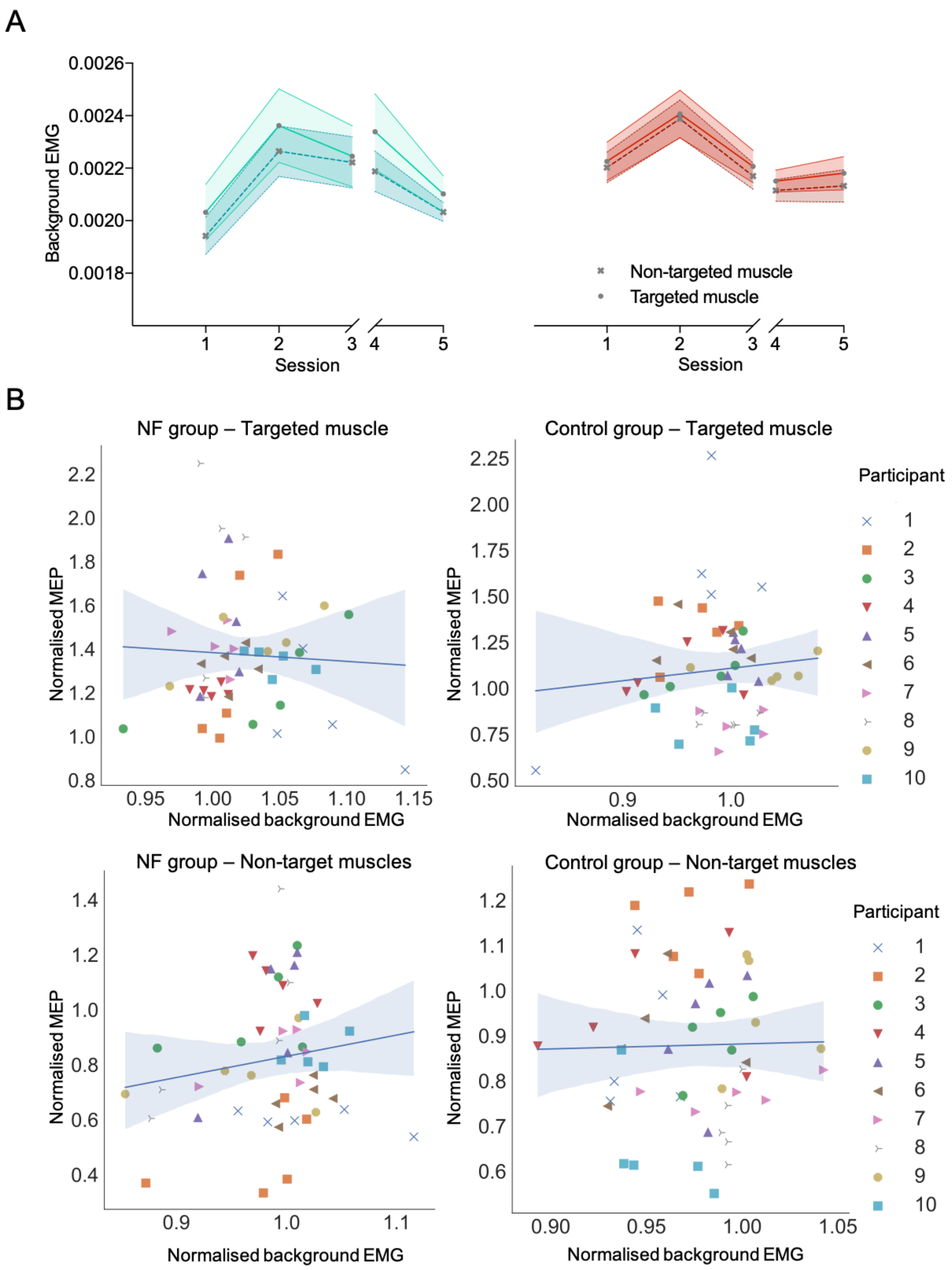


***Supplementary figure 2***. *A) We performed mixed effects analyses on the background EMG (bcgEMG) data to investigate any potential influence bcgEMG could have had on our main MEP results. Both the 3-way (group x session x target muscle) and the 2-way (group x target muscle) interactions were not significant ($X^2(4,10)=0.756$, $p=0.94$; $X^2(1,10)=0.001$, $p=0.99$),*

*thus we conclude participants in the NF group did not utilise bcgEMG to support their performance. B) We have additionally correlated the normalised bcgEMG activity with the corresponding MEP activity for the targeted and the non-targeted muscles separately. None of the correlations were significant (NF group, Targeted muscle ρ=0.121, p=0.404; NF group, Non-targeted muscles ρ=0.102, p=0.482; Control group, Targeted muscle ρ=-0.003, p=0.982; Control group, Non-targeted muscles ρ=0.053, p=0.716).*

Supplementary figure 3

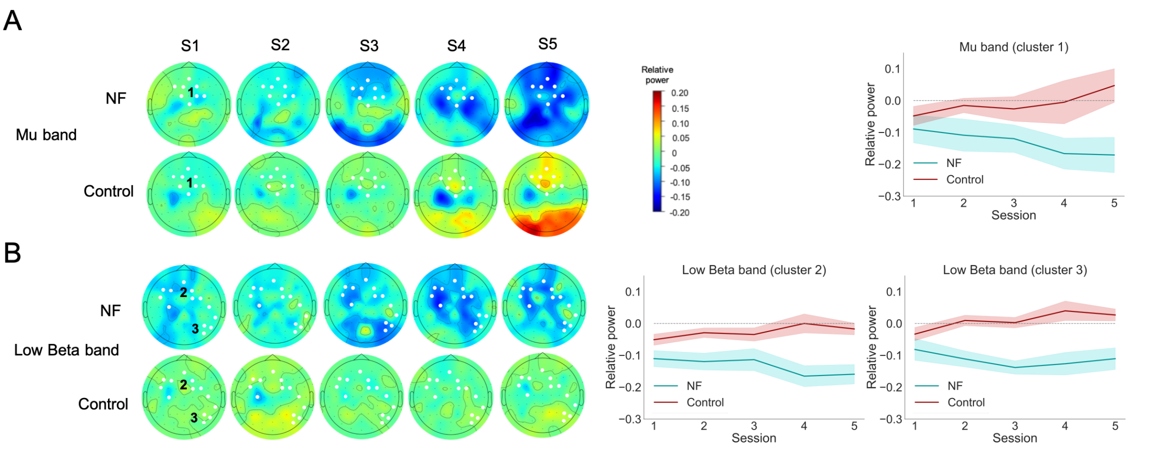


***Supplementary figure 3.*** *Non-corrected clusters A) The topoplots show changes in relative power in the averaged 2 – 3.5 s of MI period for the mu band. There is an increasing ERD trend visible across sessions in the NF group while the Control group shows smaller ERD changes and some event related synchronization (ERS). The marked electrodes formulate a cluster showing an interaction (session x group) in relative power changes, as identified by cluster analysis. To visualize this trend the activity over the 8 electrodes formulating the cluster was averaged and plotted across sessions and between groups. B) The bottom panel illustrates the same analysis performed for the low beta frequency band. ERD activity is larger and more distributed in the NF group. Two separate marked clusters suggest the session x group interaction trend, the frontocentral (cluster 2) and the parieto-occipital (cluster 3). The average activity of each cluster is plotted separately across sessions and between groups.*

***Supplementary table 1**. Since our findings are mostly communicated using chi-square ($X^2$) tests we have additionally calculated achieved power based on the achieved effect sizes and degrees of freedom. Average achieved power, on all the tests where we report a significant value, was 0.87.*

| Effect size w | Degrees of freedom | Achieved power |
|---|---|---|
| 1.6 | 14 | 0.93 |
| 1.34 | 8 | 0.88 |
| .67 | 5 | 0.13 |
| 1 | 1 | 0.88 |
| 1.98 | 1 | 0.99 |
| 1.17 | 4 | 0.85 |
| 0.9 | 2 | 0.72 |
| 0.74 | 1 | 0.63 |
| 1.09 | 4 | 0.79 |
| 1.2 | 4 | 0.87 |
| 1.03 | 1 | 0.9 |
| 1.56 | 1 | 0.99 |
| 1.62 | 1 | 0.99 |
| 0.54 | 1 | 0.4 |
| 0.54 | 1 | 0.4 |
| 0.75 | 1 | 0.65 |
| 0.42 | 1 | 0.26 |
| 3.3 | 1 | 1 |

***Supplementary table 2**. List of motor imagery strategies suggested to participants prior to the start of the task. Partially adopted from Majid et al. (2015).*

| **Upregulation strategies** |
|---|
| Focus your attention on the cued finger by thinking about moving it. |
| Imagine performing a functional movement (playing a piano or a keyboard) |
| Think about tensing the cued finger and keeping it tensed. |

| |
|---|
| Imagine moving your finger in different directions. |
| Think about isolating the cued finger. |
| Use any other strategy you come up with. |

| **Downregulation strategies** |
|---|
| Divert your attention away from the cued finger by thinking about moving the other. |
| Divert your attention to another part of the body. |
| Think about tensing the cued finger so it remains still. |
| Think about that muscle going numb or going on ice. |
| Think about suddenly stopping a movement of the cued finger. |
| Use any other strategy you come up with. |
